# Biological communities as interacting compartments: thermodynamic properties and diversity indices

**DOI:** 10.1101/188813

**Authors:** Fernando Meloni, Gilberto M. Nakamura, Alexandre Souto Martinez

## Abstract

Diversity indices provide simple and powerful metrics for assessing biological communities. Based on entropy measures, the approach considers statistical and thermodynamic inferences to deduce ecological patterns. However, concerns exist regarding the accuracy of diversity indices. Because relative quantities depend on the sorting of organisms *(e.g.*, guilds and species) and their interactions, field observations carry inherent imprecision, thus leading to misinterpretation. Here, we present a framework that is able to appropriately achieve the thermodynamic properties in ecological systems and ensure the inference power. We demonstrate that effective abundances rather than raw abundances provide a trustful estimator of probabilities, which is evaluated through massive tests. We use empirical and synthetic data to show the advantages and reliability of this new framework under a broad range of conditions. The tests demonstrate that the replication principle is always optimized by the new estimator. Compared to other methods, this approach is simpler and reduces the importance of schemes used for sorting organisms. We highlight the robustness and the valor of effective abundances for ecological contexts: *i)* to assess and monitor the biodiversity, *ii)* to define the best sorting of organisms according to maximum entropy principles, and *iii)* to link local to regional diversity *(α-, β-*, and *γ*-diversity).

## I. INTRODUCTION

The assessment of biodiversity is a primary concern among ecologists. They are interested in monitoring species and ecosystems to explain how climate, soil type and several other environmental features affect organisms and their organization in nature^15,50^. Beyond basic knowledge, the motivation for this interest has increased over the past decades due to the increasing human impacts on climate and natural ecosystems that place species survival and ecosystem services at risk^2,26^.

The best appraisal of patterns in biological communities considers details of several taxa, such as their biology, genetic variability, ecology, behavior, and so forth. However, this level of information is rarely available for practical contexts, and objective measures are considered along with external inferences to deduce the community patterns^54,55^. Taxonomic/functional compositions, the shape of the curves describing species-abundance distributions (SAD), and the ecological indices consist of distinct approaches for measuring and comparing biological communities^50^. Each approach addresses levels of information, and their use shows advantages and disadvantages for distinct contexts ^41,55^. Among these metrics, ecological indices deserve special attention, and we explore them in this paper.

Also referred to as diversity indices, ecological indices are deduced from the Boltzmann-Shannon-Gibbs entropy, and they are proposed to measure order-disorder in biological communities. These indices enjoy advantages from well-supported concepts, interpretations, formulations, and inferences from information theory, physics, statistics, and thermodynamics, and they allow deducing further relationships from data^13,15,16,36,74,78^. Briefly, the calculation of diversity indices considers the number of categories and the number of organisms (abundances) as primary information ^50^. The *W* categories are defined by any criteria of interest *(e.g.*, species, genera, behavior, guild, and so forth) and are used for sorting the organisms, while relative quantities provide a quantitative inference. This type of basic information can be obtained for a broad range of practical conditions and reflects the applicability of diversity indices. Furthermore, diversity indices provide reliable information about global patterns of biological organization even if information about organisms is limited.

Accordingly, once the *W* categories are determined *a priori*, the abundance *A*_*k*_ considers the number of organisms in the *k-*th category and determines the probability *p*_*k*_ = *A*_*k*_/*A*, for 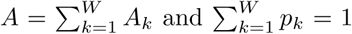. These probabilities are used to calculate the classic Shannon diversity index *H* ^50^ as follows:

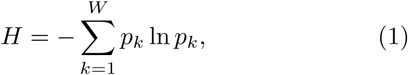

which is equivalent to the Shannon entropy ^74^. In practical contexts, the *H* values of biological communities living under distinct influences are compared by similar sampling effort. In such cases, the variation *∆H* is inter-preted as a metric that addresses the influence of a particular driver *(e.g.*, climate changes or human impacts) on the ecological framework ^13,50,53^.

In addition to the *H* index, there are several other ecological indices that provide complementary aspects of the biological communities. The notation

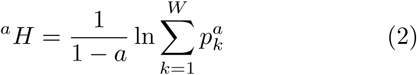

unifies the three most important ecological indices^13,15,36^: the species richness *W* (number of categories), the *H* index (Shannon entropy), and the Ginni-Simpson index 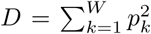, where the last infers the curve asymmetry of ranked relative quantities. For *a* = 0 and *R* ≥ 2, we have _0_*H = ln(W)*, which is the harmonic mean; for *a* → 1, ^1^*H = H*, which is the geometric mean; and for *a* = 2, ^2^*H* = 1/*D*, which is the arithmetic mean. These cases are special cases of the Rényi generalized entropy and are important because they predict useful relationships. One expects to find correlations between the abundances of organisms and the values assumed by distinct diversity indices, such as ^*a*^*f(A)* ≡ ^*a*^*H*, which also implies that *g*(^0^*H*) ≡ ^1^*H* and so on ^15,36^. These correlations match the concept of the replication principle in ecological contexts^13,15,38^, and they are true only if the additive and extensive thermodynamic properties are ensured^13,34,78^. Assuming this circumstance, the entropy observed for small samples can be re-scaled to predict the entropy of the entire system. This type of inference would be used, e.g., to explain how the biological diversity is spatially partitioned (*γ*-, *α*- and *β*- diversities)^12,38^. Although *H* and other diversity indices provide an interesting approach to assess and interpret biological communities, tests using empirical data often show a different reality, and the topic remains open ^10,12,14,24^.

The inconsistency between theoretical predictions and empirical results is justified by the lack of a precise definition of appropriate classes for ecological systems^34^. A common approach for sorting organisms considers taxonomic arguments, where individuals are strictly grouped by their phylogeny. This method is intuitive but leads to granular schemes that poorly represent ecological constraints. In fact, taxonomic sorting reflects the evolutionary history of groups better than the immediate effect of environmental forces on niche occupancy ^72^, often resulting in probabilities *p*_*k*_ that fail in evaluating the diversity patterns ^76^. In contrast to taxonomic schemes, the guild approaches employ functional traits and few taxonomic arguments, maximizing the niche context for sorting organisms. The individuals are grouped because they share similar traits and explore similar niches, a scheme that minimizes ecological redundancies. Consequently, guild methods tend to improve the assessment of environmental disturbances on biological communities 9,37,43,72. Multiple arguments (such as phylogeny, morphological traits, genetic information, behavior, niche occupancy and *ad hoc* information) can be addressed together to elaborate constraints and produce distinct *a priori* hypotheses, and they are compared next in accordance with MaxEnt principles ^13,34,35^. The arrangement that better achieves the thermodynamic properties defines the most plausible granular scheme. However, the approach also brings important concerns. For instance, knowledge of tens, hundreds, or sometimes thousands of distinct organisms is a prerequisite for accurate results using guilds. Beyond the necessity of specialists, this implies some subjectivity in selecting the traits and respective weights (if weights are addressed). Finally, guilds are considered for particular groups of organisms and under specific situations *(e.g.*, tree species in forests, arthropods in soil, fishes in lakes, and so on), and they cannot be readily inferred for wide proposes. Therefore, the opinion and context may affect the evaluation of probabilities *p*_*k*_, the values assumed by the diversity indices, and the subsequent conclusions.

Notwithstanding the importance of the assortment of organisms, we emphasize here that the granular scheme is only one side of the problem for the accuracy for biodiversity assessment. We claim that ecologists should address further sources of data variability before defining the best assortment. Ecological systems admit complex spatial-temporal dynamics (many of which are poorly understood), which finally drives the presence and quantities of organisms. Beyond environmental influences, such dynamics derive from intricate food webs and interaction networks, which finally frame the biological organization in the ecosystems. The consequence is a mutual and non-trivial dependence of relative quantities. Because ecological information is generally acquired through field observations and samplings, fluctuations in relative abundances may accrue relevant “noise” in empirical data^45,51^, leading to uncertain probabilities *p*_*k*_ for diversity indices. Consequently, the reproducibility of ecological experiments remains a topic of scientific concern^7^.

Taking these arguments into consideration, we conjecture that insights taken from non-equilibrium systems ^19,40,52,80^ could be adapted to a static approach for reducing the effects of short-range fluctuations on observed quantities. In this paper, we explore the inherent inaccuracy of diversity indices as a consequence of a poor interpretation of raw abundances when used for estimate probabilities. We propose a new estimator that is able to separate data variations produced by entropic and non-entropic drivers, assuming samples as stationary (or as *quasi-stationary)* states. The approach is demonstrated to optimize the results obtained by diversity indices, making them reliable tools for assessing biological organization. The additive and extensive thermodynamic properties of biological communities are recovered from effective abundances, thus allowing a broad range of inferences about time, space, and external influences since the adequate context is considered. Extensive tests with empirical and synthetic data remove any doubt regarding the applicability and reliability of the new estimator of probabilities. This new estimator of probabilities represents a shift in our understanding about abundances, which is taken as the result of interacting compartments rather than independent ones. Our rationale, formulation and interpretations broadly match the framework presented by Würtz and Annila^80^, with differences for the general interpretation and mainly for practical application. We discuss the phenomenological process behind the quantitative changes in the data structure, as well as the potential of this approach to answer classic problems concerning biodiversity patterns.

The remainder of this paper is organized as follows. In Sec. II, we present the rationale used to obtain an alternative estimator of probabilities, which enables optimizing the thermodynamic properties in ecological systems. In Sec. III, we use data of ground arthropods sampled in a successional gradient to show that organism sorting schemes are insufficient for assessing the entropy production in ecological systems, elucidating how effective abundances provide a distinct approach to improve the biodiversity assessment. In the same section, we test the framework in further empirical data, showing that thermodynamic properties are recovered in distinct communities, including plants, arthropods, mammals, fishes and others, that may used to infer regional diversity from few samples and is minimally affected by the shape of SAD. We finally discuss the implications of our findings and the phenomenological aspects concerning the topic in Sec. IV.

## II. ENTROPY OF BIOLOGICAL COMMUNITIES AND ESTIMATORS OF PROBABILITIES

Contextualizing the problem, empirical data are generally obtained from limited samples, and the dynamics driving the spatio-temporal patterns of diversity under *in situ* conditions are hardly accessible. As distinct influences on relative abundances, on the one hand, we have the phenomena that alter the free energy in ecosystems and directly modify the ecological framework *(e.g.*, vegetation suppression, ecological succession, climate changes, and so forth). These phenomena disturb ecological systems and lead to a system reorganization, a consequence of the second law of thermodynamics. We name these phenomena entropic influences, or *φ*. On the other hand, predation, competition, seasonality, and so forth may also affect the quantities of organisms, but these are non-entropic influences, or *9*. At homeostasis or any stationary state, such phenomena make abundances fluctuate (regularly or not) over time around a fix point. Despite being thermally neutral, these fluctuations address systematic sub-and/or overestimation of organisms in samples. Consequently, the non-entropic influences impart a high level of uncertainty to *A*_*k*_ values (primarily because of the sampling process) and hinder the assessment of entropy variation. Separating entropic from non-entropic effects on observed abundances is possibly the most important barrier for assessing the effects of environmental changes on biological diversity^11,17,25,32,42,44,71,73,79^

In their theoretical study, Würtz and Annila^80^ considered the paths to solve this question. The authors considered ecosystems as systems out of equilibrium, and they interpreted the ecological interactions as chemical potentials. Using ecological succession as context, the authors explicitly deduced the time and matched the thermodynamic entropy to ecological succession. However, the most important insight was the estimator of probabilities, which is assumed as the result of a product. Unfortunately, this topic has remained poorly explored until now. We develop an independent reasoning that widely converges to a similar mathematical framework. Because the concept was previously proven, we spend less time with justifications in order to focus on the applicability and implications. Our approach also has some differences because we address each sample as a stationary state, a static approach that does not demand that time be explicitly considered ^20,21^. In fact, the stationarity addressed here is a condition imposed to the sample, and it is not a necessary condition for the system. The approach strictly focuses on variability in abundances, which for methodological and practical concerns enables wider inference power from *∆H* because time, space or any other driver can be deduced from appropriate experimental designs. With some contextualization, we next present a step by step description of the alternative probability estimator *q*_*k*_, which is a function of *A*_*k*_ and *φ* but not necessarily proportional to them.

Following Haegeman and Loreau^33^, the Shannon entropy [Eq. (1)] describes the organization of biological communities in a given ecological system. The main aspect that is necessary to obtain a reliable value of *H* is the proper definition of groups according to which the organisms are classified. As previously mentioned, there are several ways to group organisms in compartments, and we start with the taxonomic view. At a basic granular level, species define the categories, and the abundance *A*_*k*_ represents the number of organisms in the *k*-th species of the system, while the total abundance is 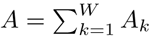 and *W* is the species richness. The ratio

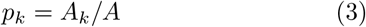

estimates the occurrence probability of finding a given element of the *k-th* compartment. Naturally, this defini-tion ensures that 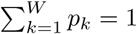.

Despite the intuitive formulation of *p*_*k*_, applying Eq. 3 to Eq. 1 raises several experimental questions. For instance, small changes in abundance *∆A*_*k*_ = *ε_k_A* with *ε*_*k*_ ≪ 1, either due to sampling errors or measurements in different ecosystems, produce corrections 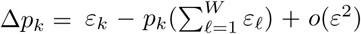. Since *ε*_*k*_ are assumed to be small, corrections to the estimators themselves are also very small. However, they may produce large corrections to the entropy,

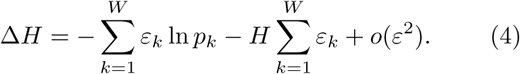

For clarity, let us consider the particular case of a single variation in abundance, i.e., *ε*_*k*_ ≡ *ε*. Under this circumstance, 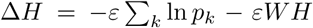. Note that in the first parcel, *εΣ*_*k*_ *ln p*_*k*_, compartments with small probability *p*_*k*_ ≪ 1 provide considerably larger contributions to *∆H* than compartments with a larger presence in the ecosystem. This result should be compared against compartments with equal probability *p*_*k*_ = 1/*W*, ∆*H* = *εWH* - *εWH* = 0. Therefore, large deviations in *H* arise due to the estimator Eq. (3), making the index sensitive to population dynamics.

Another pressing concern regarding the reliability of Eq. (3) as an ecological index revolves around the organism interactions and their sorting in categories. These interactions are expected to shape the various levels of the biological organization either directly or indirectly. In thermodynamics, even though interactions are known *a priori*, the sheer amount of components in the system prohibits the exact computation of properties. The chemical potential *μ* overcomes this problem by introducing an effective energy change caused by the addition of another particle to the system, including temperature and pressure. In this way, the chemical potential encodes all resulting effects of underlying interactions in the system, with Gibbs free energy as *G = μN*, for *N* identical particles. For ecological problems, the translation is straightforward: we associate a chemical potential *μ*_*k*_ = *μ*_*k*_(*φA*_1_,…, *A*_*W*_) to the *k*-th compartment such that the total Gibbs free energy is

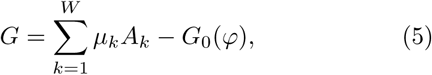

where *G*_0_(*φ*) depends only on entropic variable *φ*. From Eq. (5), it is clear that the chemical potentials *μ*_*k*_ represent weights to the compartment occupation *A*_*k*_, from which we infer the estimator

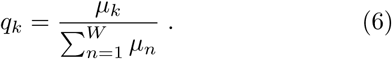

One reobtains the estimator *p*_*k*_ when *μ*_*k*_ ∞ *A*_*k*_, *i.e.*, the effort required to add another organism to the *k-*th compartment increases linearly with *A*_*k*_.

For more complex environments, we should define or estimate the values *μ*_*k*_ in some other way. Indeed, from the Maxwell relation *β*_*μ*_*k*__ = -(*∂H*/*∂A*_*k*_) and 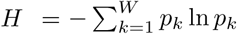 (Eq. (1)) with estimators *p*_*k*_ = *A*_*k*_/*A* 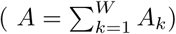, one obtains

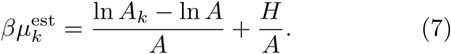

It is easy to show that the term ln(*A*_*k*_) - ln(*A*) in Eq. (6) refers to the *H*_*max*_ (maximum entropy possible) only if all relative abundances are the same (Suppl. 1). For ecological systems, further considerations are necessary. The ratio *H*/*A* ≪ 1 for biological communities since *H* ~ *o*(1), whereas *A* ≥ *o*(10^3^). Moreover, *A*^−1^ ln *A* is a constant for fixed *A* and only shifts the free energy. From these considerations, we have

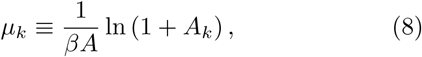

which in turn produces our estimator corresponding to the *k-*th compartment

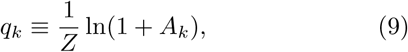

with the partition function 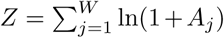 guaranteeing that 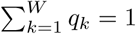. Comparing distinct estimators and their thermodynamic interpretation, the Gibbs partition function is

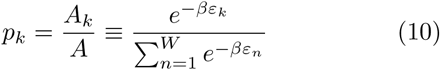

while *q*_*k*_ is

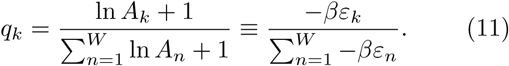

The estimator *q*_*k*_ vanishes for vanishing *A*_*k*_ similar to *p*_*k*_. However, *q*_*k*_ exhibits notable differences from *p*_*k*_ in terms of the relative variations 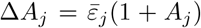 with 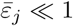,1

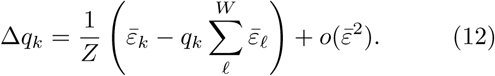

A brief comparison between Eq. (12) and *∆p*_*k*_ reveals an interesting fact. Let us assume that both estimators *q*_*k*_ and *p*_*k*_ share similar numerical values, *p*_*k*_ ≈ *q*_*k*_, with constant 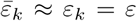. These particular constraints are unlikely to naturally occur in uncontrolled environments; they serve only to highlight general properties. For instance, despite the numerical equivalence between the estimators in this hypothetical scenario, the variations *∆q*_*k*_ are reduced by a constant factor *Z* ~ *o(W)* > 1, which mitigates fluctuations of abundances *A*_*k*_. For more realistic systems, we consider the fluctuations *∆φ* due to entropic influences:

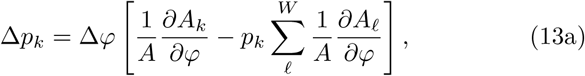

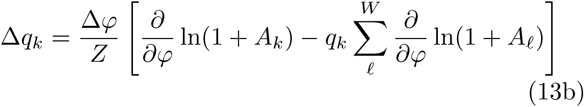

Other than the factor *Z*^−1^, ∆*q*_*k*_ and *∆p*_*k*_ differ mostly in the way the variations behave: *∆q*_*k*_ depends only on relative variations, whereas *∆p*_*k*_ requires the variation of *A*_*k*_ with *φ* in addition to the total value *A* = *A*(*φ*). Consequently, *q*_*k*_ may produce a better separation of effects from the different compartments in the system when compared to *p*_*k*_ for the same compartmentalization scheme. In other words, the estimator *q*_*k*_ reduces the interdependences among the various compartments.

From the relation *q*_*k*_ ≤ *A*_*k*_/*Z*, one obtains the inequality *q*_*k*_/*p*_*k*_ ≤ *(A/Z)*. Therefore, the positive index

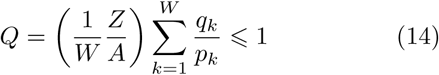

which measures the deviation between the probability estimator *q*_*k*_ from the more common estimator *p*_*k*_. This assumption shifts the concept of units in ecological systems, meaning that only exponential variations in relative abundances are properly inferred in terms of entropy variations. Note that our rationale matches the general MaxEnt framework for thermodynamic variational principles^19-21^, whereas our partition function *Z* is similar to that assumed in Würtz and Annila^80^. In fact, Eq. 11 demands deduction and inferences, and we present them in terms of abundances. Eq. 5 indicates that abundances are the result of distinct terms. To clarify this topic, we consider *F*_*k*_(*φ*) to be the entropic influence on the abundance of the *k-*th compartment, and *θ*_*k*_ is the non-entropic influence of the *W* compartments on quantities found in the *k*-th compartment.Then, any entropy production is related to *∆F*(*φ*), while non-entropic effects result from the internal dependence among categories (biological interactions, population dynamics, and so forth).

Applying these terms to abundances, the classic approach assumes that

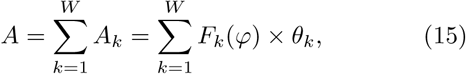

where *A* is the result of an arithmetic relation among the *W* compartments, a framework that makes little sense in terms of dependent compartments. For instance, consider the effects produced by the introduction of an exotic species to a random ecosystem. One does not expect that the addition of *n* predators will reduce exactly *n* prey, as assumed by the arithmetic ratio. In fact, a predator consumes several prey, affecting the growth rate of prey because the number of prey also affects the growth rate of predators. Therefore, this mechanism produces a cascade of influences on relative abundances, which are better expressed by multiple dependences^80^. Considering *A* as the result of a product rather than a sum, Eq. 15 assumes that

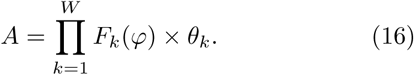

The continuous observation during the stationary state would imply that *θ*_*k*_ is a function able to describe how *A*_*k*_ fluctuates around the fixed point determined by *F*_*k*_(*φ*). However, the sampling process makes *θ*_*k*_ assume discrete values indicating whether *A*_*k*_ deviates from the value produced by *F*_*k*_(*φ*) at the right moment of the sample. Then, *θ*_*k*_ = [*λ*_*k*_*F*(*φ*)]/[*F*(*φ*)], where *λ*_*k*_, *∀k* is a Lagrange multiplier, and

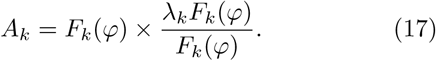

For *θ*_*k*_ = 1, the influence of the further compartments on the *k^th^* is null, whereas for *θ*_*k*_ ≠ 1, some influence is detectable.

As categories interact, the quantities in one category are expected to increase at the expense of a decrease in another during the stationary state, maintaining the thermodynamic balance according to the second law. Consequently, all values assumed by *θ* alternate around 1 through the *W* compartments. Concerning Eq. 16, entropic and non-entropic influences on abundances *A*_*k*_ can be isolated at the logarithm scale as

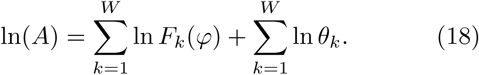

For the consequence of 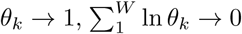, and therefore, we have the “effective” abundances *Â*_*k*_ as

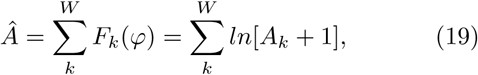

where unit summed to *A*_*k*_ guarantees the relation even when one or more categories are not represented in the sample. This framework states that *A* is a function of “observed” raw abundances *A*, and it exclusively holds the entropic influences on *A*_*k*_. Therefore, it is expected that

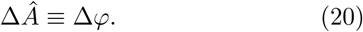

Importantly, as *θ*_*k*_ → 1, *∀k*, the average value 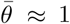, and the expression 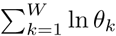 can be rewritten as *W*[*ln*(*W*^−1^], making Eq. 18 assume that

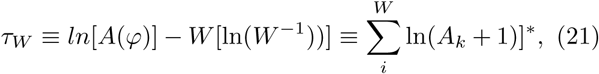

where *τ*_*W*_ is the optimized entropy for the granular scheme that produced *W*. The errors associated with Eqs. 19 - 21 reflect whether the assumption *θ*_*k*_ → 1 deviates from the true distribution produced by *θ*_1_,…, *θ*_*W*_, inferring whether the assumed granular scheme deviates from the actual maximum entropy possible for the system. Accordingly, the multiplicative approach and consequent probabilities calculated as *q*_*k*_ offer a robust approach for comparing granular schemes and for assessing the entropy production in ecological systems. Because time is not necessary and thermodynamic properties are recovered, further inferences can be explored, such as the space and consequently the partitioning diversity. In the following, we shall test the plausibility of this rationale using empirical and synthetic data.

## III. TESTING THE FRAMEWORK IN EMPIRICAL DATA

### Entropy production in soil systems

In the first test with empirical data, we verify whether the aforementioned framework is applicable for practical contexts. For this purpose, we use TGA, which is a data set that contains information of plant biomass and counts of ground arthropods in a successional gradient of tropical forests. Concurrent to the granular scheme, we aim to evaluate how diversity indices of ground arthropods respond to the entropy production when the classic probabilities, *p*_*k*_, and the new one, *q*_*k*_, are committed.

TGA covers 13 forests in distinct stages of development through a gradient constituted by 10 reforestations and 3 forest remnants. All sites are located in Southeast Brazil and bear a high diversity of tree species (Atlantic forests). Data of plant biomass are removed to infer the entropic influences on the diversity of ground arthropods. Then, samples of tree densities (*ind./m*^2^), area of trunks (*m*^2^), and litter biomass (*kg/m*^2^) are converted to only one variable, *φ*, by using principal component analysis^45,59^. *φ* represents a general quantitative descriptor of forest development, and its non-dimensional values stand for the linear coefficients of the eigenvector of the largest eigenvalue (see Supplement 2 for details). Concerning ground arthropods, the input data are counts of ground arthropods sampled in soil and litter during winter and summer, totaling 40 samples per site (see Supplement 2 for details). Part of these data were previously analyzed to explain the colonization of ground arthropods during the development of the forest ^57,60^. The organisms are sorted by species (occasionally morpho-species), and data of distinct samples are joined in only one large sample per site. With data of fauna in hand, we consider the species sorting to calculate the number of categories *W* (species richness), the abundance *A* and the classic Shannon diversity *H* (Eq. 1) for the 13 sites of TGA. Next, we plot respective values against *φ* quantities, and we observe the correlation between them.

Figures 1A-C depict that the variations in the three metrics, namely, *W, A* and *H*, show no correlation with the values of *φ*. Consequently, two possible explanations may be assumed. The first one admits no cause-effect relationship between the forest development and the biological organization of ground arthropods. This explanation would imply that the successional process does not produce entropy in litter and soil layers or that ground arthropods are not sensitive to such a process. This hypothesis appears to be implausible due to strong experimental evidence attesting to the contrary^1,4,17,18,27,57,60,64^. The alternative explanation assumes that the cause-effect relationship exists, but *W, A* and *H* are unable to properly assess entropy production in soil communities. We assume that the second explanation is correct because it is corroborated with experimental evidence.

**Figure 1.**
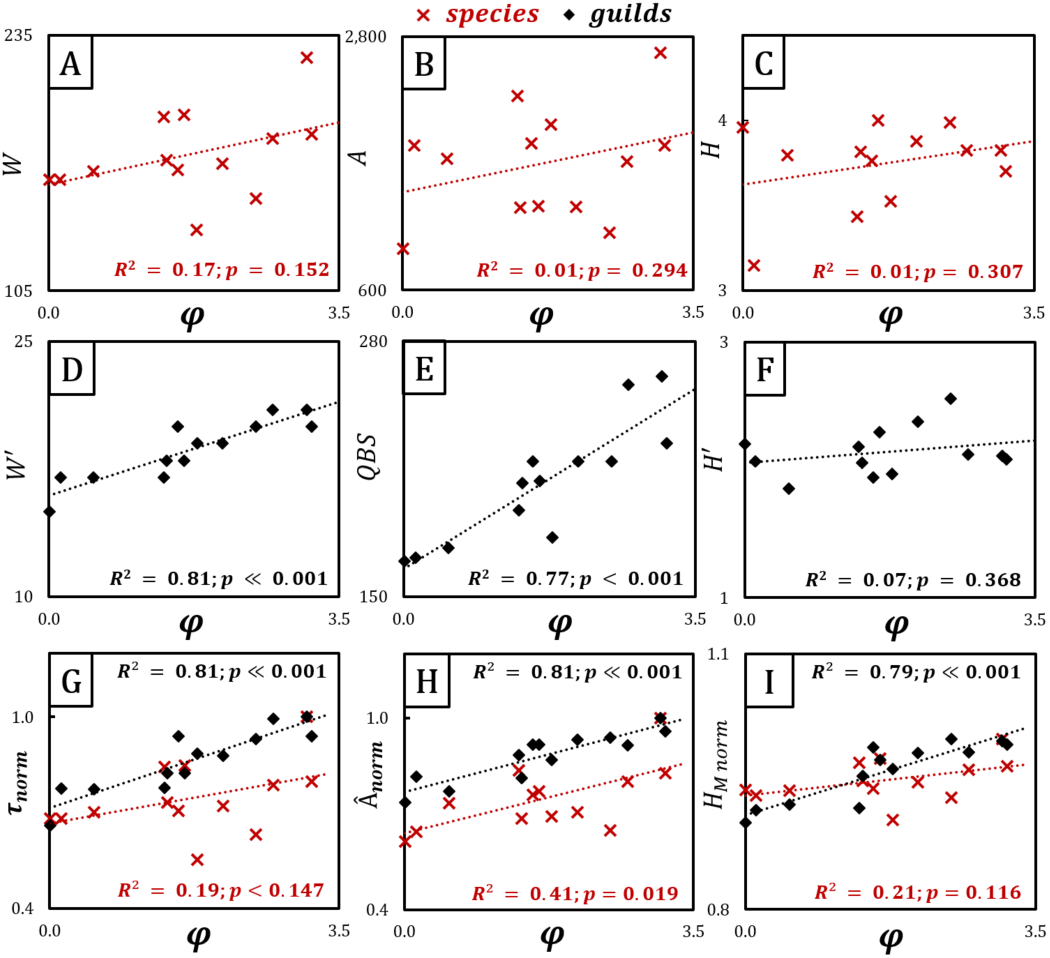
Relationship between successional process and ecological indices of ground arthropods considering a gradient of thirteen sites *(n* = 13). *φ:* indicator of forest development, obtained from a PCA conducted using values of tree dominance, tree density and litter biomass. For ground arthropods, *R:* species richness; *A:* total abundance; *H:* Shannon diversity index using species as compartments; *W:* number of guilds according to Parisi *et al.^67^; QBS:* Quali-tat Biologica del Suolo, an indicator of soil health based on ground arthropods^67^; *H’:* Shannon index obtained for guilds as compartments; *τW = ln(A)* - *W ln(W^−1^*, the maximum entropy; 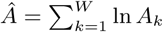, the effective abundance; and *HM*: Shannon entropy achieved by probabilities *q*_*k*_ = ln*(A_K_)/Â*. A-C show that *R, A* and *H* of ground arthropods are poorly correlated with the forest development. The use of guilds increases the correlation between the number of categories *W* and the successional stage, as does *QBS*. However, the use of guilds as compartments produces values of entropy *H’* that are poorly correlated with the forest development. Finally, the use of *q*_*k*_ probabilities, derived from the effective abundances approach, show that the forest development affects the organization of ground arthropods, mainly when *q*_*k*_ is considered together with guilds.

There is a clever way to avoid the concerns around *H* and related indices: sorting organisms with categories other than taxonomy, in which relevant biological and ecological constraints are explicitly addressed. In fact, conceptual and empirical evidence suggest that guild approaches may improve the extraction of ecological information because they fairly reflect the niche occupancy _1_,_9_,_61_,_67_ Ideally, functional approaches would pursue new categories that alter *H* values and improve the thermodynamic properties. Consequently, a better inference about entropy production for the TGA case is also expected, and we test this hypothesis.

Thus, we sort the arthropods of TGA in guilds following the criteria described in Parisi *et al*.^67^. Tailored from ground arthropod data, this methodology collects informative content about the biological quality of the soil and produces a reliable indicator of soil health, the *QBS (Qualità Biological del Suolo)* _5_,_30_,_47_,_61_,_67_. Accordingly, arthropods are sorted by high-level taxa and morphological traits into taxonomic-functional categories (≈ 28). Each category receives an *ad hoc* score, the EMI (eco-morphological index), that provides a quantitative inference of how adapted its organisms are to soil niches (Supplement 3). For a random sample, the EMI scores of all represented categories are summed to produce the *QBS* value. Note that relative abundances show limited importance for the calculation of *QBS*, and single or multiple occurrences at the same *k*-th category result in equal contributions. Concerning the TGA example, one expects that the number of EMI categories and *QBS* values increase over the forest development as a consequence of soil maturation^17^. We then conjecture that the EMI categories could also improve the assessment of entropy production in TGA if abundances are used rather than the *ad hoc* values.

In the following, we refer to the EMI categories as guilds, and we add the prime symbol to differentiate its use (primed) from the use of species (unprimed). For arthropods sorted according to the EMI categories, *W* corresponds to the number of guilds, and *QBS* is the respective index calculated following the approach of Parisi *et al.*^67^. Using guilds as a granular scheme, we also calculate *H’*, as *p’*_*k*_ = *A’*_*k*_/*A*, where *A’*_*k*_ is the relative abundance of the *k-*th guild. Then, we plot *W’, QBS*, and *H’* against the *φ* values, and we compare these results with those obtained for species.

Figures 1D-E depict how the number of guilds *W’* and the *QBS* index vary as functions of *φ*. The linear fits indicate that both quantities correlate with the forest development, inferring that the successional process affects the organization of soil communities. In fact, guilds join several organisms in a few categories, which naturally reduces the total variance. However, the same result does not hold for *H’* (Figure 1F). In comparison to *H* (species), *H’* reveals that guilds produce negligible improvements in the assessment of entropy production. Furthermore, the *H’* behavior is in clear contrast to that of *QBS*, although both metrics address the same granular scheme. How is this result possible? Because *QBS* computes *ad hoc* quantities and *H’* computes abundances, it is clear that the estimator *p*_*k*_ makes *H’* inefficient. Therefore, the granular schemes alone cannot assess the entropy production in systems such as TGA, making the use of an appropriate estimator of probabilities necessary.

Then, we follow the framework presented in Sec. II, considering abundances as the result of interacting categories. Employing *q*_*k*_ probabilities, we calculate the Shannon index (Eq. 1) for two granular schemes: species and guilds. We use the notations *H*_*M*_ (species) and *H’*_*M*_ (guilds) to differentiate the indices calculated by *q*_*k*_ from those calculated by *p*_*k*_. The indices *τ* and *Â*, predicted by Eq. 19 and Eq. 21, are also calculated, and all them are then plotted against *φ*.

Figure 1G shows that correlations between *τ*_*W*_ or *τ’*_*W*_ with *φ* are compatible with the respective correlations observed for *W* and *W’*, which confirms the gain produced by guilds in comparison to the species scheme (Figures 1A,D,F). Furthermore, the tests indicate that the *q*_*k*_ probabilities always attain better inference about *φ* than the *p*_*k*_ (Figures 1H-I). For both species and guilds, the effective abundances *A* are correlated with the values of *φ*, with even the granular schemes still being important. Once TGA addresses successional context, guilds naturally produce stronger inference power about entropy production than species. When *q*_*k*_ is employed together with guilds, abundances finally reveal the entropy production in the TGA data set, with *Â’* achieving even better results than *QBS* (Figure 1E,H). As predicted by Eq. 19, *H’*_*M*_ also correlates with *φ*, which is evidence that effective abundances allied to guild compartments recover the thermodynamic properties in the TGA case. Such optimization is predicted by Eq. 21, and since it is confirmed, it ensures the replication principle. Because the replication principle is of great importance for thermodynamic inference, we apply further tests to explicitly evaluate this hypothesis.

### Recovering the replication principle in soil systems

The replication principle states that diversity indices are functions of abundances^13,15^, which requires that extensive and additive thermodynamic properties be ensured. Concerning the extensive property, it is expected that an increase of abundance leads to a monotonic increase in the number of compartments and in entropy values. For the additive property, it is expected that the entropies of samples considered separately are equivalent to the entropies of samples considered together, implying the independence of categories. Subsequently, we evaluate the reliability of such relationships in the TGA example.

Objectively, we test whether *q*_*k*_ can actually optimize the extensive property in the TGA case by evaluating whether the increase of *Â* leads to a monotonic increase in *W, H*_*M*_ and *τ*_*W*_. Such correlations are expected only when the replication principle is ensured. We also test the influence of granular schemes by evaluating the curves obtained for species and guilds separately. As predicted in Sec. II, Figure 2 depicts that the values of *H*_*M*_ and *H’*_*M*_ linearly correlate with respective effective abundances *Â* and *Â’*, the number of categories *W* and *W’*, and maximum entropy *τW* and *τ’*_*W*_. These results explain the better inferences about entropy production depicted in Figure 1, suggesting that the *q*_*k*_ probabilities clearly achieve the extensive properties in the TGA case. Furthermore, the results clearly indicate that the optimization of the extensive property does not necessarily depend on the granular scheme, which also confirms the predictions of Sec. II.

**Figure 2.**
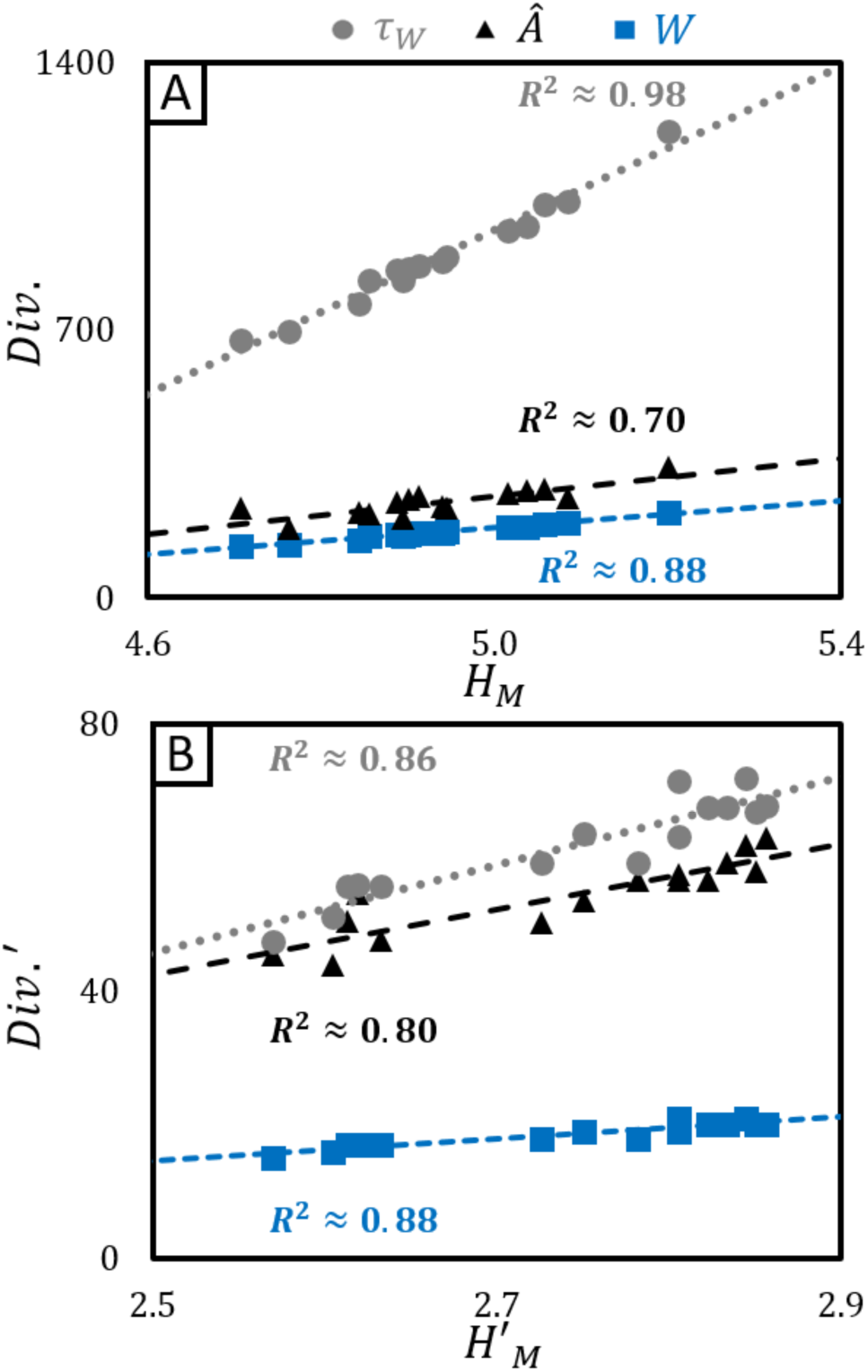
Thermodynamic relations for ecological indices of ground arthropods sampled in secondary tropical forests when using probabilities *q*_*k*_. **A**: arthropods sorted as species; **B**: arthropods sorted as guilds^67^. *Div.:* diversity indices; *W:* number of categories; *Â:* effective abundances; *τ*_*W*_: maximum entropy for *W* compartments and *A*. The figure shows that probabilities *q*_*k*_ can recover the extensive and additive properties of ground arthropod communities. Under such circumstances, the thermodynamic relationships are guaranteed, thus allowing the prediction of how changes in the number of organisms are related to the number of compartments, as well as the system organization.

Next, we explicitly evaluate whether the additive property is also recovered by *q*_*k*_ in TGA. The additive property states that *HA + HB* ≡ *HAB*, where *A* and *B* are samples, while *AB* is their union. This would be final evidence that *q*_*k*_ ensures the replication principle (thermodynamic properties) in ecological systems. To avoid any influence of the entropy production on TGA data, which address correlations among samples, we develop a strategic test to properly evaluate the additivity in this data set.

The test consists of a shuffling routine followed by systematic comparisons between the variances (*σ*^2^) of sample data separately with the variances of the same sample data together.The sites of TGA (*m* = 13) are labeled as *U*_*a*_, *U*_*b*_,…,*U*_*m*_ and are used to create *c* clusters *C*_1_,…,*C*_*c*_. Each cluster combines arthropod counts of four random sites, such as *C*_*1*_ = (*U*_*a*_ ∪ *U*_*b*_ ∪ *U*_*c*_ ∪ *U*_*d*_), *C*_2_ = (*U*_*a*_ ∪ *U*_*c*_ ∪ *U*_*d*_ ∪ *U*_*e*_), and so on. Next, a second set of *x* clusters is created, now combining the clusters *C*_1_,…, *C*_*m*_ two by two, such as *C*_(1,2)_,…, *C*_(*x*-1, *x*)_, where *C*_*i,j*_ = (*C*_*i*_ ∪ *C*_*j*_). With this shuffled structure in hand, we calculate the variances (*σ*^2^) of respective probabilities, such as 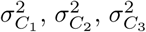, and so on, for the first group of clusters, and 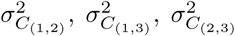, and so forth for the second one. Because the additive property explicitly concerns the independence of compartments, the granular scheme is of great importance. Then, we propose four distinct treatments to evaluate the subject:

- Treatment 1: *W* for categories and *p*_*k*_ for probabilities;
- Treatment 2: *W* for categories and *q*_*k*_ for probabilities;
- Treatment 3: *W’* for categories and *p*_*k*_ for probabilities;
- Treatment 4: *W’* for categories and *q*_*k*_ for probabilities.

Briefly, each treatment evaluates the linear correlation between 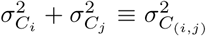. A good linear fit indicates that the treatment allows achieving the additive property. Conversely, the lack of a correlation suggests that the thermodynamic property is poorly achieved. Figures 3A-D depict the results.

**Figure 3.**
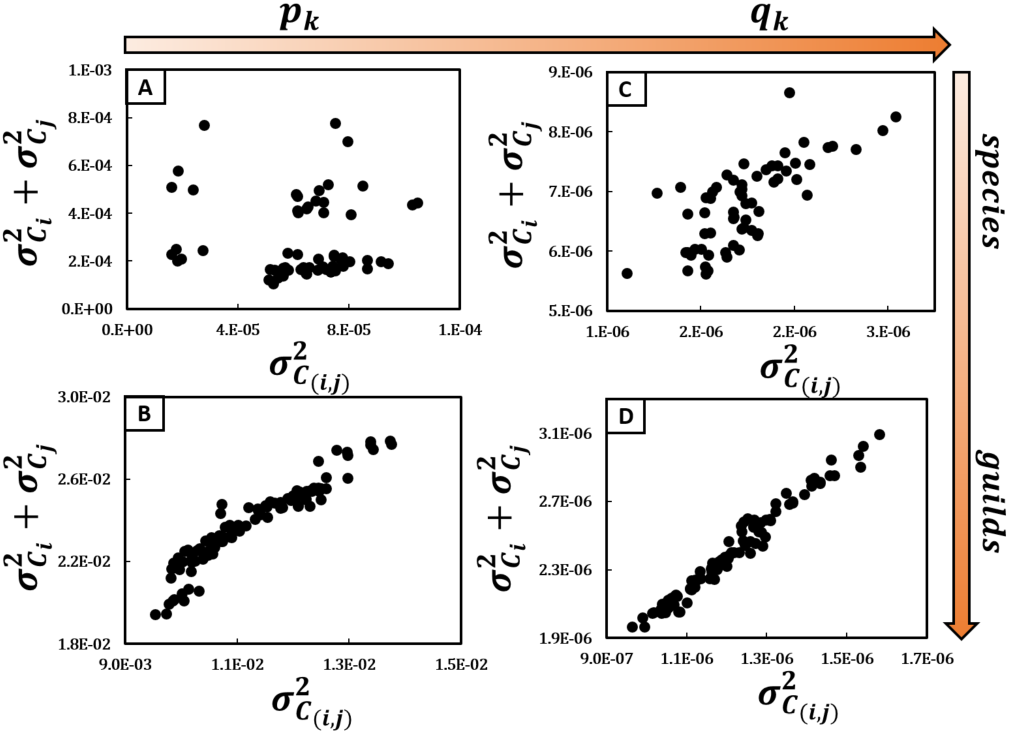
Comparing the variances *(σ*^2^) of probabilities obtained using distinct approaches for quantities of ground arthropods found in secondary Atlantic forests. Thirteen sites showing different levels of development are used to create multiple arrangements (n=63). **A**: species compartments and *p*_*k*_; the sum of variances shows a poor linear correlation with the variance of sums (*R*^2^ < 0.01; *p* = 0.593), indicating that additive and extensive properties of the systems are not achieved. **B**: guilds^67^ and *p*_*k*_ (*R*^2^ ≈ 0.90; *p* ≪ 0.001); **C**: species and *q*_*k*_ (*R*^2^ ≈ 0.55; *p* < 0.001). In both cases, the correlation between the sum of variances and the variance of sums was improved in comparison to A, suggesting that the thermodynamic properties are better assessed. **D**: guilds and *q*_*k*_; the excellent linear fit (*R*^2^ ≈ 0.98, *p* ≪ 0.001) indicates that the additive and extensive properties of the ecological system are recovered by addressing both approaches together.

Figure 3A indicates that, for Treatment 1, the sum of variances is not compatible with the variance of distinct communities together. Therefore, the species scheme and *p*_*k*_ probabilities cannot ensure the additive property. Figures 3B and 3C indicate that Treatments 2 and 3 partially recover the additive property, implying that the granular scheme and the estimator of probabilities consist of distinct approaches to reduce data variability in ecological systems. Finally, Treatment 4 (Figure 3D) shows that the guild scheme and *q*_*k*_ probabilities employed together produce an excellent linear fit between 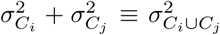, indicating that the additive property is completely recovered.

Once more, the tests support the results depicted in Figure 1 and the predictions of Sec. II. Therefore, the tests confirm the independent effect of *q*_*k*_ probabilities on the optimization of thermodynamic properties of soil systems. Although the *q*_*k*_ probabilities ensure the replication principle in the TGA case, an inference about general ecological systems requires further tests. Thus, we subsequently evaluate the reliability of the approach in further contexts.

### Beyond soil systems: the generality of the result obtained

Notwithstanding the predictions of Sec. II and evidence from soil systems, we hypothesize that the optimization of thermodynamic properties by *q*_*k*_ cannot hold for broader ecological contexts. Hence, we evaluate the replication principle in general situations, taking ten data sets available on-line and calculating the entropy produced by the estimators *p*_*k*_ and *q*_*k*_. These data sets concern a broad range of biological communities in worldwide ecosystems, including mammals, perennial plants and ants from US deserts^3,62^; fishes and benthic invertebrates from temperate lakes^48,49^; ground arthropods sampled in Italy’s organic and conventional crops^61^; butterflies living in open areas from Colorado, US^63^; plant species growing in steppes from Oklahoma^56^; understory plant communities of temperate forests from Canada^66^; and tree communities of tropical forests from India^69^. The experimental conditions also greatly vary among studies. There are times series, simple samples in spatially explicit conditions, counting of individuals, biomass estimation, frequencies, and so forth. We provide a list of the main characteristics of these data sets in Supplement 3, but we encourage a search in the original papers for more details (all of them are available as open source data).

This broad range of experimental conditions is suitable for an important propose: assessing and comparing the replication principle and the thermodynamic inference produced by the classic and the new estimators of probabilities. Note that we are not attempting to replicate the original experiments using a new tool, or evaluating the reliability of the previous findings. In fact, several details about the original experimental designs and inferences about external influences are not addressed here, because they are pointless for our propose. Accordingly, climate, human manipulation, entropy production, granular schemes, and other aspects described in these data sets are here considered as only sources of entropy variation. Our only intention is to check the consistency of the estimator *q*_*k*_ concerning the replication principle, observing how the diversity indices fit variations in organism quantities. Based on the consistency of thermodynamic laws, it is expected that the data variability and the distinct conditions considered by these data sources minimally affect the general patterns and that the abundance variability present in each data set would provide sufficient information to observe the thermodynamic properties. Finally, we aim to evaluate the applicability of the framework described in Sec. II as a general tool for assessing biological diversity in ecological systems.

Accordingly, for a data set *S* containing *n* samples, we simply calculate the diversity indices of each sample, and we analyze the correlation among distinct indices under the conditions found in *S*. Because we intend to prove the optimization of the replication principle, we evaluate the correlation between the same diversity indices considered by Hill numbers ^13,15,36^, although not necessarily as Eq. 2. For *p*_*k*_, we observe the values of *W*, *H* and *D* as functions of *A*, whereas for *q*_*k*_, we observe *W*, *H*_*M*_ and *D*_*M*_ as functions of *Â’*. Furthermore, we assess the correlation between *W* and *H* and *H*_*M*_. In contrast to the tests with TGA, only one granular scheme is assessed per data set. Based on MaxEnt principles, the best estimator is considered to be the one capable of producing index values that optimize the inference produced by relative quantities. The results are depicted in Figure 4 and Figure 5.

**Figure 4.**
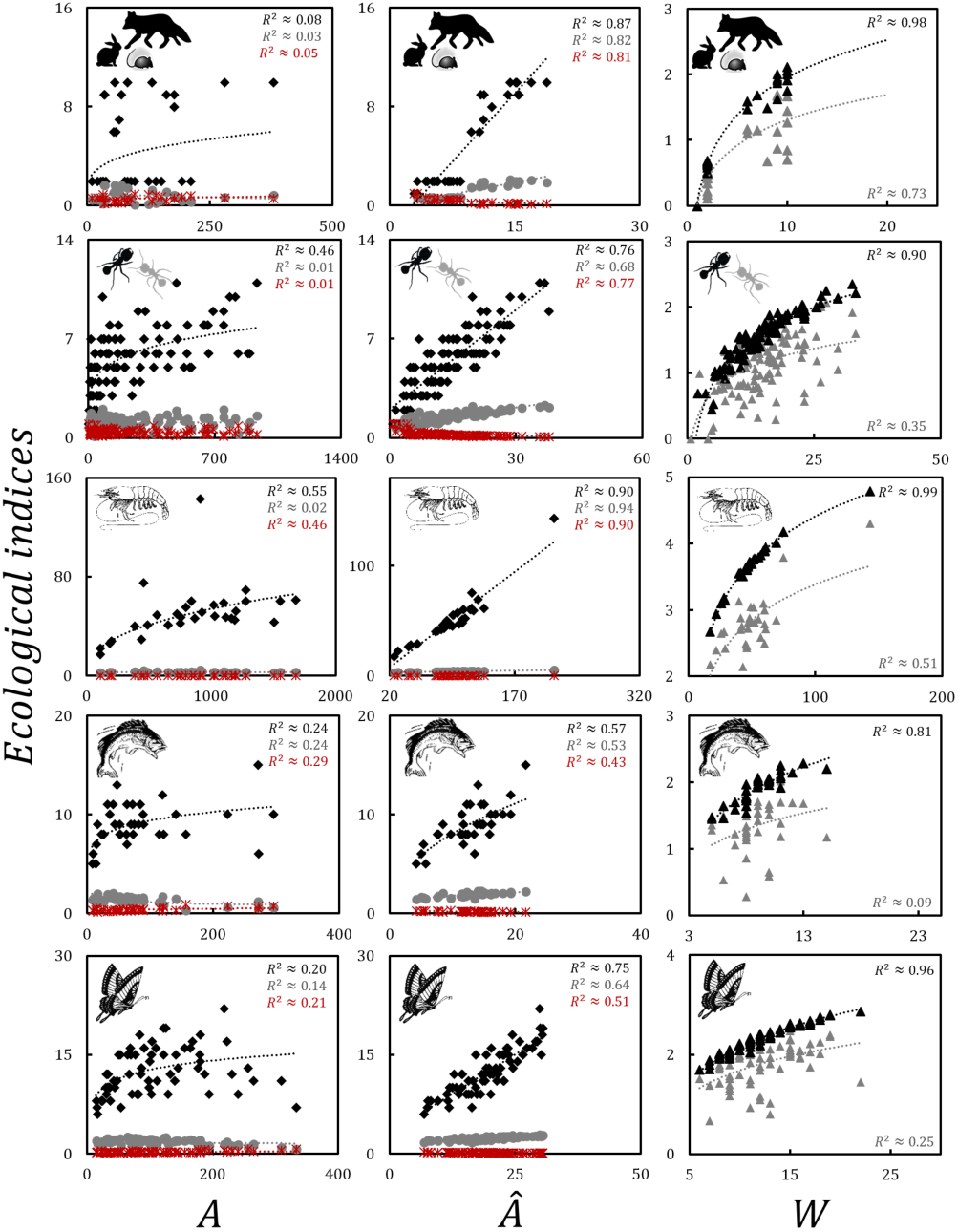
Relationship between quantities of organisms and entropy measures according to distinct estimators of probabilities. *A:* abundances; *Â*: effective abundance; *W:* number of compartments (species, genera, guilds, and so forth); black diamonds represent the number of compartments *W;* gray diamonds represent the Shannon entropy; and dark red asterisk represents the Gini-Simpson asymmetry *D*. On the left, the indices are calculated for probabilities taken as *p*_*k*_ = *A*_*k*_/*A*, and in the middle, by probabilities as 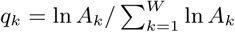. On the right, black triangle represents the *H* obtained by *q*_*k*_, while gray triangle represents the *H* obtained by *p*_*k*_. Accordingly, probabilities *q*_*k*_ improve the maximum entropy for distinct biological communities, revealing their additive and extensive properties. In lines: 1^*st*^: Mammals from Utah, US^3^; 2^*nd*^: Ants from Chihuahuan Desert, Arizona, taken by baits^62^, *n* = 135, *W* = 20; 3^*rd*^: Benthic invertebrates from North Lake region^48^, *n* = 30, *W* = 19; 4^*th*^: fishes from temperate lakes of North America taken by electro fishing^49^, *n* = 35, *W* = 27; 5^*th*^: Butterflies from open areas of Colorado, US,^63^ *n* = 66, *W* = 58. **Art**: <u>*oogazone.com*</u>.

**Figure 5.**
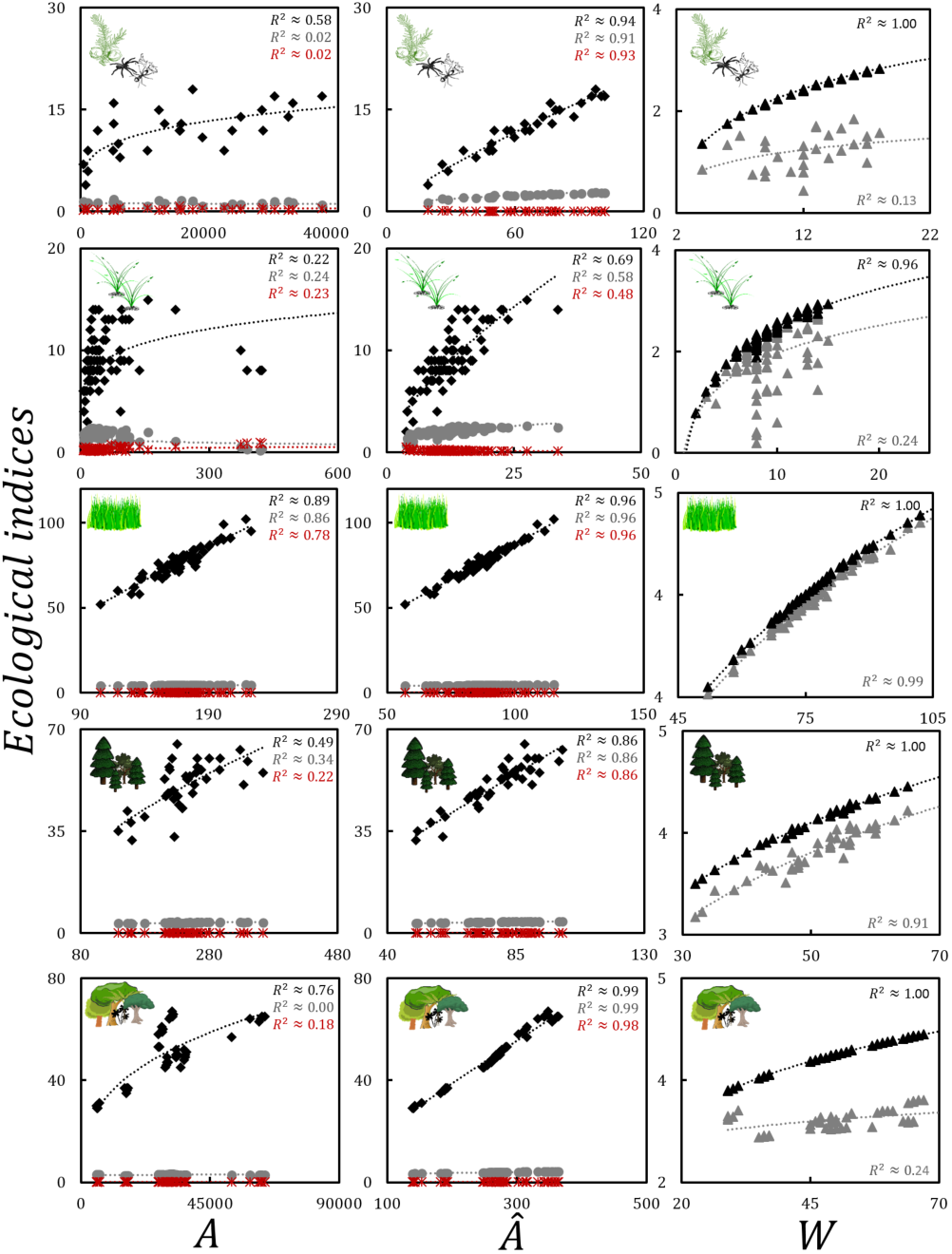
Relationship between quantities of organisms and entropy measures according to distinct estimators of probabilities. *A:* abundances; *Â*: effective abundance; *W:* number of compartments (species, genera, guilds, and so forth); black diamonds represent the number of compartments *W;* gray diamonds represent the Shannon entropy; and dark red asterisk represents the Gini-Simpson asymmetry *D*. On the left, the indices are calculated for probabilities taken as *p*_*k*_ = *A*_*k*_/*A*, and in the middle by probabilities as 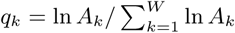. On the right, black triangle represents the *H* obtained by *q*_*k*_, while gray triangle represents the *H* obtained by *p*_*k*_. Accordingly, probabilities *q*_*k*_ improve the maximum entropy for distinct biological communities, revealing their additive and extensive properties. In lines: 1^*st*^: Abundance of soil arthropod groups found in cultivated soils from Italy *n* = 28, *W* = 28^61^; 2^*nd*^: Perennial plants from Chihuahuan Desert, Arizona, *n* = 42, *W* = 59^62^; 3^*rd*^: Vegetation cover of grass and shrub species in prairie steppes from Oklahoma, US^56^, *n* = 60, *W* = 320; 4^*th*^: Semi-quantitative crown cover from trees in the Lac Croche temperate forest, Quebec, Canada, *n* = 40, *W* = 72^66^; 5^*th*^: Basal area of trees in tropical forests from India, *n* = 46, *W* = 100. **Art:** <u>*oogazone.com*</u>.

Concerning the ten data sets, the use of *p*_*k*_ almost always produces values for the diversity indices (*W*, *H*, and *D*) that poorly correlate with abundances *A*. Even the correlation between *W* and *H* is important only for one case (Figure 5^56^). Therefore, the test corroborates the poor inference provided by the classic estimator of probabilities, which assumes *a priori* the biological categories as independent compartments. Conversely, probabilities *q*_*k*_ always achieve strong correlations between effective abundances and the ecological indices or between *W* and *H*_*M*_ (Figures 4 and 5). Note that probabilities *q*_*k*_ are appropriate even for the case in which *p*_*k*_ probabilities are satisfactory.

Therefore, the results corroborate the evidence provided by the TGA example, as well as the prediction of Sec. II. Concerning the broad range of conditions regarded by this test, the results depicted in Figures 4 and 5 stress the valor of the framework presented here for the assessment of diversity patterns in ecological systems. These new and strong evidence reinforce that effective abundance, as well as the entire conceptual framework behind its assumption, consists of a trustful manner to recover the thermodynamic entropy of ecological systems, which is a substantial gain in the inference power produced by diversity indices based on Boltzmann-Gibb-Shannon entropy. In fact, the approach is so consistent in all experimental situations that we further proceed to find additional inferences, as follows.

### Probabilities *q*_*k*_ as a tool to link the partitioning of diversity

The examples presented above provide strong evidence supporting the use of *q*_*k*_ probabilities to achieve the thermodynamic properties in ecological systems. In turn, we conjecture that *q*_*k*_ probabilities can also be employed to solve an old question in ecology: the partitioning of biodiversity^24^. Briefly, the problem consists of whether the diversity of local samples represent the entire diversity found in the region. The local diversity is named *α-*diversity, the regional diversity is named *γ*-diversity, and their relationship is molded by the *β*-diversity. This important subject has generated debate among ecologists for decades and still lacks a consensus, with dozens of frameworks already proposed^24^. Because probabilities *q*_*k*_ optimize the extensive properties in general ecological systems, we conjecture that an appropriate experimental design could reveal how diversity patterns are affected by the sampling area. If the correlation between sampling area and *H*_*M*_ is confirmed, then the *q*_*k*_ probabilities would provide the basis for definitively solving the dilemma, at least under the thermodynamic perspective.

Hence, we use a data set that contains counts of vascular plants, where the samples are taken in an experimental design with spatial correlation^65^. This data set considers a large parcel of 256*m* × 256*m*, partitioned into 256 samples of 16*m* × 16*m* arranged side by side, as in a square lattice. We calculate *A, Â, W, H*_*max*_, *H* and *H*_*M*_ for one parcel alone, then for two parcels together, and then for three parcels together, and so on, until all 256 of the parcels are addressed together. We plot the respective values against the respective area (*m*^2^) represented by the data. For extensive systems, a monotonic increase of all indices is expected. Figure 6 shows that the abundances, effective abundances and the number of species monotonically increase as a function of the sampling area, as expected for extensive systems. The same holds for *H*_*max*_ and for *H*_*M*_, but not for *H*. In fact, the entropy calculated by *p*_*k*_ produces an anomalous behavior, suggesting that entropy would decrease despite the increase in sampling area. Therefore, no doubts exist that *q*_*k*_ can actually recover the extensive properties of ecological systems and can be used to link *γ*-diversity to *α*-diversity only using a scale correction, while *p*_*k*_ is not a proper metric. Naturally *β*-diversity is a function that links local and regional diversity and it can unequivocally predict how much of the regional diversity is represented in a set of samples.

**Figure 6.**
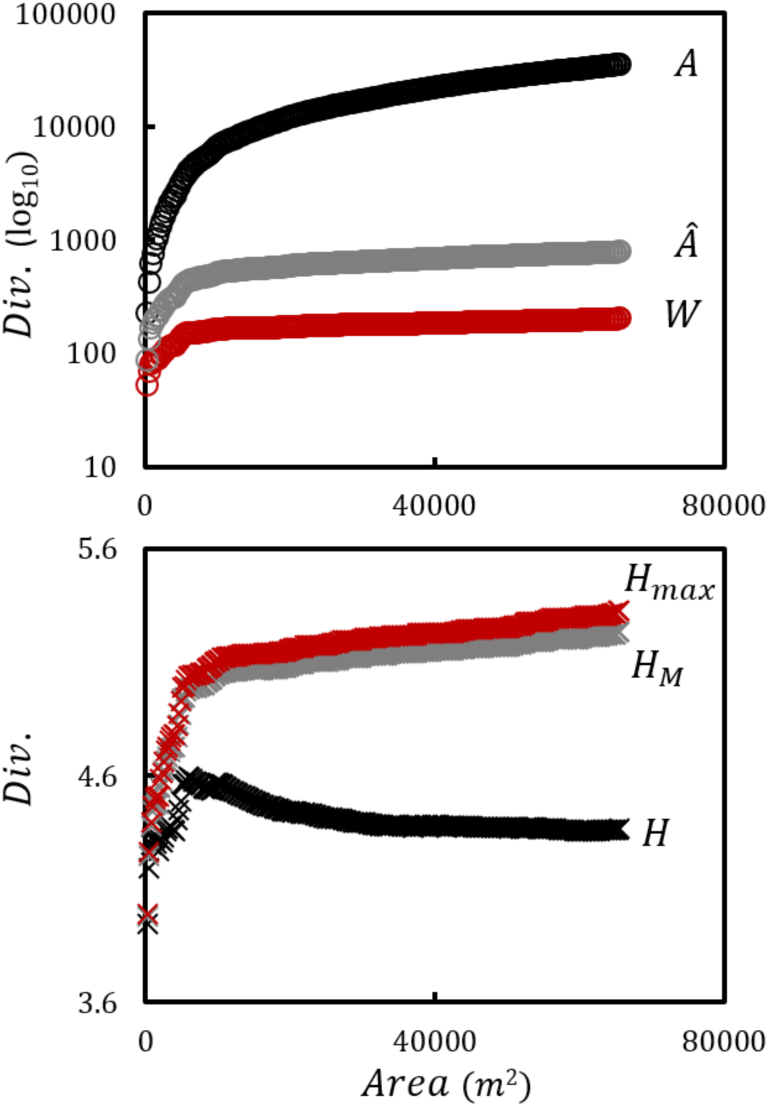
Influence of sampling area (*m*^2^) on the diversity of plants according to different estimators of probabilities. *Div.:* diversity metrics; *A:* abundances; *Â*: effective abundance; *W:* species richness; *H*_*max*_: maximum entropy possible for *W* compartments; *H:* Shannon entropy calculated by *p*_*k*_ = *A*_*k*_/*A*; *H*_*M*_: Shannon entropy calculated by 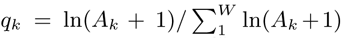. The analyses consider the data of Palmer *et al.*^65^. On the top, the increase in sampling area led to the monotonic increase of abundances, effective abundances and species richness. The bottom shows that *H*_*max*_ and the entropy produced by *q*_*k*_ (*H*_*M*_) monotonically increase with the area, whereas the entropy produced by *p*_*k*_ (*H*) presents an unpredicted pattern. Therefore, only *H*_*M*_ achieves the extensive properties of this system.

### Underlying ecological mechanism

The last doubt regards a possible influence of underlying ecological mechanisms on the inference produced by *q*_*k*_. In fact, the forces driving the shape of SADs are poorly known, and the method presented here makes no *a priori* inferences about them. In fact, distinct mechanisms can produce SADs with different shapes, which could carry consequences for the assessment of entropy in biological communities^68^. Consequently we test whether *q*_*k*_ probabilities are sensitive to the SAD shape using synthetic data.

We employ computational algorithms to generate data distributed as four probability density functions (PDFs) 31,70 for our treatments, including *a)* Poisson, *b)* exponential, *c)* lognormal, and *d)* power law. In each case, we produce *n* = 50 data vectors 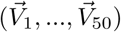, where 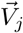 represents a “pseudo” biological community found in the *j^th^* site. The length of 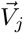 denotes *W*_*j*_, while its data enters the relative abundances *A*_1_, *A*_2_,…, *A*_*W*_*j*__. In all treatments, we intentionally produce “pseudo” communities, where respective species richness *W*_*j*_ varies according to a Poisson distribution (∧ = 35). Furthermore, we monotonically increase the parameter values of generator functions, guaranteeing that all the treatments contain some entropy variability (see Supplement 4 for details). Similar to the tests with empirical data, we compare the correlation between diversity indices produced by *p*_*k*_ and *q*_*k*_ in accordance with thermodynamical predictions.

Briefly the results obtained for synthetic data (Figures S1-S4) show that probabilities calculated as *p*_*k*_ achieve extensive properties for systems whose SADs are distributed as a Poisson or exponential PDF. However, *p*_*k*_ was hardly efficient for SAD distributed as lognormal or power law functions. Conversely probabilities calculated as *q*_*k*_ optimized the thermodynamic properties in all treatments. Therefore, the use of *q*_*k*_ probabilities is demonstrated to be robust even when data variability is large, as in lognormal and power law cases, and the optimization of entropy is hardly sensitive to the underlying mechanisms driving the SADs.

### IV. DISCUSSION

We present an important advancement for the assessment and monitoring of biodiversity. In fact, our framework does not develop a new entropy or create a new diversity index. Rather, one simply employs a more appropriate estimate of abundances to well-known concepts and formulations. Then, the novelty concerns which quantities can actually recover the thermodynamic properties of ecological systems. The several tests presented here indicate that our approach maximizes the replication principle, providing a more precise measure of order-disorder patterns of biological diversity. The potential update in the ecological discussion is substantial because several themes showing conflicts between theoretical predictions and empirical data can now be reviewed without further samplings. Next, we discuss some particular implications.

The most important aspect of our findings concerns the estimator of probabilities. The approach proposed here is highly logical in biological terms. The quantitative dependence among categories is a well-known aspect of any ecological system that has remained poorly understood until now. Then, the major conceptual shift occurs for the calculation of probabilities and quantitative interpretations. The dependence among compartments assumes multiplicative processes rather than additive ones and yields important consequences for diversity indices and their inference power. For instance, only exponential increases in the number of organisms (or biomass) produce detectable variations in terms of entropy. Second, the emergence of a new category in ecological systems *(e.g.*, a new guild) depends on the geometric increase of the entire system. Such deductions appear to explain the general interpretation of biomass pyramids^77^, for which distinct trophic levels show a geometric dependence. Further evidence comes from the effects produced by the inclusion/exclusion of species in natural ecosystems, such as in biological invasions. Such events can affect the entire community in non-trivial manners, which cannot be explained by the simple additive processes^22,46^.

Our tests demonstrate that the classic estimator *p*_*k*_ that relies on additive processes yields weak results when assessing diversity in empirical data. The inefficacy of *p*_*k*_ is not new in ecological discussions, but the consequences of its inaccuracy have not received suitable attention. We wonder how many of the conflicting aspects concerning the biological diversity are in fact consequences of misinterpretations produced by this problematic estimator. In our tests, we include some important topics that exemplify our concern: the thermodynamic inference provided by diversity indices, the maximum entropy produced by distinct granular schemes, the assessment of entropy production in changing ecosystems, and the explicit relationship between local and regional diversities, which leads to the partitioning of diversity. All these topics represent a large part of the long-term discussion in ecology, and the estimator *q*_*k*_ appears to offer a potential solution for several of them. Therefore, we claim that the estimator *p*_*k*_ should be definitively left aside in favor of the adoption of *q*_*k*_ and that the topics mentioned here be reassessed under this new approach.

The mathematical formulation proposed here follows well-established concepts and methods used to assess non-equilibrium systems^19-21^, which are adapted for a static approach. In fact, the probabilities and interpretations presented here match those proposed for ecological contexts in Ref.^80^, which surprisingly has remained poorly explored until now. In comparison with the previous findings, in which time is an inherent aspect^80^, the static approach here takes into account that each sample is being interpreted as a stationary (or *quasi*-stationary) state. Note that the hypothesis of stationarity is fragile under a more rigorous interpretation but that this poorly affects the accuracy of the calculation or the practical implications of the results. In this sense, our framework consists of an additional step in the direction of thermodynamic inferences, concepts, and views for wider ecological scopes. Once the diversity indices can produce trustworthy metrics, time, space or any other variable of interest can be experimentally contextualized and its phenomenological effects deduced from the ∆*H* obtained from samples.

Therefore, the importance of *q*_*k*_ is mostly practical and corroborated by empirical data. The framework is simple and is in accordance with thermodynamic rules, and it does not require additional theoretical assumptions and new formulations. The tests with empirical data revel how resilient the approach is to variations of granular schemes and is applicable to a broad range of ecological conditions. In fact, our set of tests could not find bounds for the use of *q*_*k*_, which is another highlight. However, the most clear advantage concerns the inference power and the prediction ability produced by the new estimator, which matches the replication principle. Because *H*_*M*_ uses a formulation similar to that used by the classic *H*, the approach shows another advantage in comparison to other recently proposed methodologies^8,16^: as previously mentioned, *H*_*M*_ can readily be employed to review previous studies without further field work. In fact, the reproducibility of results in ecological experiments is currently in question^7^, and some studies with biodiversity showing conflicting results can now be reevaluated without further data samplings.

As demonstrated, the thermo-statistical interpretation achieved by *q*_*k*_ in ecological systems provides the basis for deep advancements in the knowledge about biodiversity^12,14,24,38^. As we demonstrated here, *α*-, *γ*-, and *β*-diversities are perfectly inferred by the thermodynamic inferences. Because entropy and the number of categories scale with effective abundances, few samples can be used to estimate the regional pool of species, as well as the total and relative abundances expected for the entire region. The same is applicable to studies searching for adequate schemes for sorting organisms, as we show for the TGA example (species × guilds). Our tests are not extensive, and we argue that several other tools and methods addressing the MaxEnt principles^33-35^ can take advantage of the use of *q*_*k*_. This potential deserves particular attention in future studies.

### Phenomenological process behind the data transformation

Concerning the phenomenological implications of the new estimator, an important question arises: why does the logarithm of abundances better achieve the thermodynamic properties of biological communities? To answer this question, we need to interpret ecological patterns under a statistical view. The asymmetry of SAD curves is considered to be a universal pattern of ecological systems. Although the underlying mechanisms that produce SAD are still debatable, they are certainly related to the unbalanced fitness of species and the consequent number of respective decedents. As the logarithm function asymmetrically affects relative quantities, it is natural that log-transformed proportions are less uneven than raw proportions. This principle is well known from signal analysis, where mel frequencies^28,75^ are used to reduce the signal-to-noise ratio and provide more clear human interpretation of sound waves. In fact, humans can only distinguish great variations in sound waves, while small oscillations are imperceptible. This concept is also used in statistics, where the Box-Cox techniques^6^ are used to manipulate relative frequencies in random variables and generate Gaussian PDFs from data distributed according to another function. Similar to the TGA case, the entropic contribution for the formation of polymer chains is separated from non-entropic influences using a similar framework^29^. Therefore, it is clear that the geometric symmetry for relative proportions is not producing arbitrary results and only reflects the quantitative effects of interacting compartments on relative abundances. Therefore, the mechanism proposed here is known in other disciplines, and it is able to reduce data noise and produce a new probability distribution for a distinct physical system. In this sense, we also show its applicability for ecological systems. Compared with generalized entropy^78^, the assumptions of the use of *q*_*k*_ do not require a new granular scheme^39^ because only the quantitative signal inferred by the observables are considered.

One important aspect addressed here remained under-explored: the underlying mechanism driving the shape of the SAD curves. The shape of SAD curves and its relation with the central limit theorem^23^ can be explored to deduce the ecological dynamics. The theorem states that the additive processes would result in SAD distributed as Gaussian distributions or exponential distributions (if constrains are addressed), while multiplicative processes would result in lognormal distributions. Obviously, the theorem predicts such patterns as opposite limit cases when realizations are massive, but large spectra can be observed for limited sampling effort. Accordingly, the empirical evidence clearly shows that SAD curves are almost always distributed as a lognormal PDF, occasionally as a power-law PDF, but almost never as PDFs such as Gaussian or exponential. For the lognormal cases, we have a clear indication of multiplicative processes during the pattern formation. However, distinct processes related to sampling effort can produce a power law from data that are actually lognormal distributed^58,81^. Therefore, even for when the power-law hypothesis is the best approach to describe the SAD curves, the evidence point in the direction of multiplicative processes molding abundance, which should be explored in favor of defining the underlying mechanisms driving the biological diversity.

## ACKNOWLEDGEMENTS

We are grateful for C.R.F. Granzott’ comments during the manuscript preparation. Thanks to E. Stanley and NTL-LTER for permitting us to use their data sets. This work was supported by the São Paulo Research Foundation (FAPESP) Grant No. 2013/0G19G-4.

## Footnotes

1 Note the difference between *ε*_*k*_ = ∆*A*_*k*_/*A* and 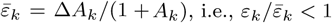

